# Anti-CD206 CAR T Cell Treatment Restores Fibrosis-Induced Loss of Dermal White Adipose Tissue

**DOI:** 10.1101/2025.10.16.682860

**Authors:** Chanhyuk Park, Asmaa O. Mohamed, Helen C. Jarnagin, Rajan Bhandari, Jason Gunn, Joana Murad-Mabaera, Noelle N. Kosarek, Fred W. Kolling, Owen M. Wilkins, Yina H. Huang, Michael L. Whitfield, Patricia A. Pioli

## Abstract

Fibrosis drives pathology in the chronic autoimmune disease systemic sclerosis (SSc), which has the highest case fatality rate of any systemic autoimmune disease with no validated biomarkers or curative treatments. Our prior work has shown that CD206^+^ macrophages and dermal fibroblasts engage in cooperative mechanisms of inflammatory and fibrotic activation in SSc. Here, we designed a targeted immunotherapeutic approach to eliminate CD206^+^ macrophages using chimeric antigen receptor (CAR) T cells. We demonstrate that systemic delivery of a single dose of anti-CD206 CAR T cells restores dermal white adipose tissue (DWAT) *in vivo*. Notably, loss of subcutaneous fat is a well-recognized but poorly understood aspect of SSc pathogenesis that precedes the development of fibrosis and is driven by changes in lineage commitment of adipose-derived stem cells (ADSCs). Using snRNA-seq and a newly-developed *in vitro* co-culture model, we report that CD206^high^ macrophages mediate ADSC shift from adipocytic to fibrotic activation in part through an IL-6-dependent mechanism. This report implicates a novel function for macrophages in the regulation of early SSc pathogenesis and is the first to establish the therapeutic efficacy of using CAR T cell immunotherapy to target macrophages in the treatment of SSc skin disease.

## Introduction

Systemic sclerosis (SSc) is a progressive, chronic multi-system disorder of unknown etiology that is characterized by immune dysfunction and fibrosis. Despite advances in disease management, including the approval of nintedanib [1] and more recently, tocilizumab, for the treatment of SSc-associated interstitial lung disease [2], the overall prognosis for SSc remains significantly worse than for other rheumatic diseases. Mortality rates are high for SSc, with 30–40% of patients dying within 10 years of diagnosis [3]. While cooperative interactions between macrophages and fibroblasts have been implicated in disease pathogenesis, the molecular mechanisms responsible for SSc development and progression are incompletely understood. To date, no therapies that specifically target pathological macrophages are in clinical use for SSc.

Inflammatory monocytes and tissue resident macrophages mediate fibroblast activation through contact-dependent and independent mechanisms. In this regard, bioinformatic analyses of tissues derived from individuals with SSc implicate profibrotic macrophages as key drivers of disease in multiple end-target organs [4, 5]. SSc macrophages, which are characterized by high surface expression of CD206, produce profibrotic cytokines and biological mediators, including TGFβ, that induce activation of fibroblasts [6]. SSc-derived fibroblasts co-cultured with SSc macrophages upregulate pathways associated with extracellular matrix (ECM) deposition and inflammation, and are characterized by increased expression of αSMA (ACTA), a marker of myofibroblasts that drive skin and lung fibrosis in SSc [6, 7]. Collectively, these studies implicate monocytes and macrophages as key effectors of skin fibrosis in SSc [8, 9], thereby linking an immune-fibrotic axis as a driver of disease.

While tissue-resident fibroblasts constitute a primary source of profibrotic myofibroblasts, many other cell types can contribute to the myofibroblast pool in SSc, including epithelial and endothelial cells, pericytes, and adipocytes [10]. Using mouse models of SSc, preadipocytes and mature adipocytes derived from dermal white adipose tissue (DWAT) have been shown to undergo adipocyte-myofibroblast transition, wherein subcutaneous adipocytes commit to the myofibroblast lineage [11, 12]. Notably, DWAT atrophy is a long-recognized clinical feature of SSc. Histological evaluation of SSc patient skin demonstrates decreased numbers of adipocytes but increased representation of infiltrating dermal macrophages, depleted DWAT and increased collagen deposition [11]. Consistent with this, loss of DWAT is associated with increased collagen deposition in multiple mouse models of fibrosis [11, 12]. Time course studies using the bleomycin-induced mouse model of SSc indicate that DWAT loss occurs early in pathology, preceding the development of skin fibrosis [12]. These results suggest loss of DWAT may play a primary or potentially inciting role in the development of dermal fibrosis. However, the cellular and molecular mediators that regulate this process are still incompletely understood.

Because our prior work implicates CD206^high^ macrophages in SSc pathology, we developed a novel therapeutic approach to eliminate these cells using chimeric antigen receptor (CAR T) cells. We now show that systemic delivery of a single dose of anti-CD206 CAR T cells restores DWAT *in vivo*. Using a newly-developed *in vitro* co-culture model, we demonstrate that profibrotic macrophages mediate the adipocytic shift to fibrotic activation in part through an IL-6-dependent mechanism. These results highlight a novel role for macrophages in the pathogenesis of SSc and show for the first time that immunotherapeutic targeting of macrophages significantly ameliorates skin disease.

## Results

### Myeloid depletion with clodronate significantly inhibits DWAT loss

Macrophages play an integral role in mediating fibrotic activation in many tissues, including heart, kidneys, and lungs (reviewed in [13]). Consistent with prior reports, delivery of bleomycin to C57BL/6 mice resulted in a significant increase in dermal thickness and attenuation of DWAT **(Figures 1A-C)** [14, 15]. To assess the contribution of myeloid cells to dermal fibrosis, clodronate was used to deplete myeloid cells in this model. As indicated in **Figure 1D**, mice were administered control or clodronate liposomes intraperitoneally three days prior to the initiation of bleomycin treatment, followed by subsequent liposome treatment every three days until the conclusion of the experiment. While dermal thickness was not significantly affected by myeloid depletion **(Figure 1D)**, mice that received clodronate liposomes had significantly thicker adipose layers compared with mice that received bleomycin alone or bleomycin plus control liposomes **(Figures 1E and 1F)**. These results implicate a role for myeloid cells in the regulation of dermal adipose loss in fibrotic skin.

**Figure 1.**
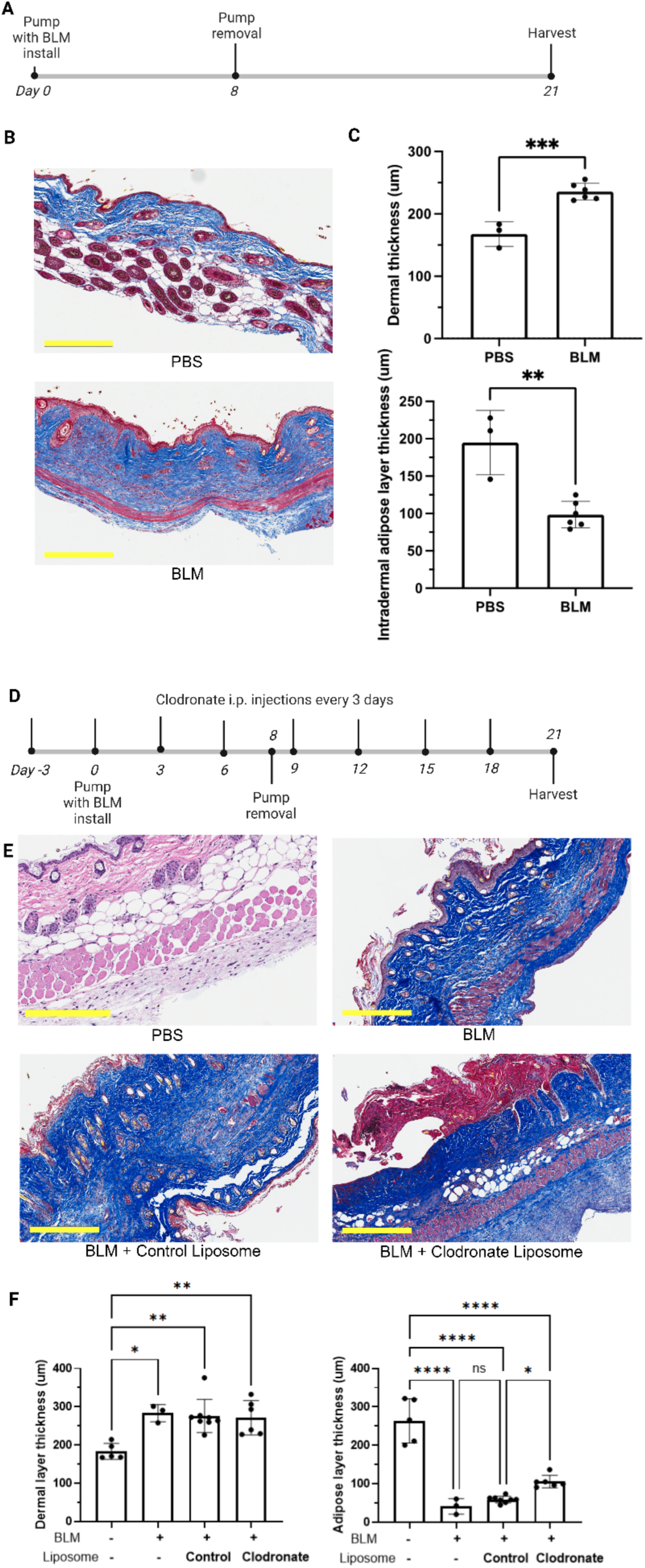
Myeloid depletion using clodronate liposomes inhibits DWAT loss in mice. (**A**) Experimental design: Osmotic minipumps with PBS or 60 U/kg bleomycin (BLM) were implanted subcutaneously on backs of 6-week-old female C57B/L6J mice at day 0, and were removed at day 7. Animals were euthanized at day 21. (**B**) Representative hematoxylin and eosin (H&E) images of PBS (top) and BLM-treated groups (bottom). Scale bar = 300μm. (**C**) Measurements of dermal and intradermal adipose tissue thickness were collected using Aperio Imagescope, where 5 measurements were collected per section. (**D**) Experimental design: Mice were treated with clodronate or control liposomes three days prior to BLM initiation, and every three days thereafter for the duration of the experiment. Animals were euthanized at day 21. (**E**) Representative dermal H&E on day 21 of naïve (top left), BLM-treated (top right), BLM and control liposome-treated (bottom left), and BLM and clodronate liposome-treated mice (bottom right). (**F**) Measurements of dermal (left) and adipose layer (right) thickness were collected as in 1C. *p≤0.05; **p≤0.01; ***p≤0.001; ****p≤0.0001.

### DWAT is significantly restored by CAR T cell-targeting of CD206^+^ macrophages

CD206 is dysregulated in a variety of pathologies, including rheumatological diseases [16]. Therapeutic targeting of CD206, which is upregulated on tumor associated macrophages, has shown significant clinical benefit in cancer [17-19], and a recent study showed that elimination of lung CD206^+^ macrophages in a bleomycin-induced mouse model significantly mitigated fibrosis without impairing host defense [20]. Notably, our prior work has shown that human SSc macrophages are profibrotic and have elevated expression of CD206 (MRC1) [6]. Given these findings and data implicating myeloid cells in SSc pathogenesis **(Figure 1)**, we designed a targeted approach to specifically interrogate the role of CD206^+^ macrophages in dermal fibrotic activation. The specificity, durability, and migratory capabilities of T cells led us to investigate the potential utility of CAR T cells in eliminating pathological dermal CD206^high^ macrophages.

We generated CAR T cells expressing a second-generation CAR construct, anti-CD206-CD8TM-CD28-CD3zeta(ζ). The design of the retroviral CAR construct was based on the Variable Heavy domain of Heavy chain (VHH) of an anti-CD206 camelid antibody (nanobody) fused with the mouse CD28 hinge, transmembrane, and intracellular domains, followed by CD3ζ signaling domain. Additionally, the CAR was cloned in *cis* with mouse truncated CD19 (tCD19) marker separated by a 2A linker **(Figure 2A)**. *In vitro* functional activity and targeted specificity of virally transduced control and anti-CD206 CAR T cells were measured by analysis of IFN-γ secretion and CD206^+^ macrophage killing. Anti-CD206 CAR T cells produced IFN-γ in a dose-dependent manner to a serial dilution of the CD206 target and recognized and killed CD206-expressing bone marrow derived macrophages (BMDMs) (**Figures 2B and 2C**). Prior to use *in vivo*, CAR T cell transduction efficiency and phenotype were assessed *in vitro*. As demonstrated in **Figure 2D**, both control and anti-CD206 CAR T cells were characterized by high viability and purity (99% CD3^+^) and were significantly enriched in CD8^+^ T cells.

**Figure 2.**
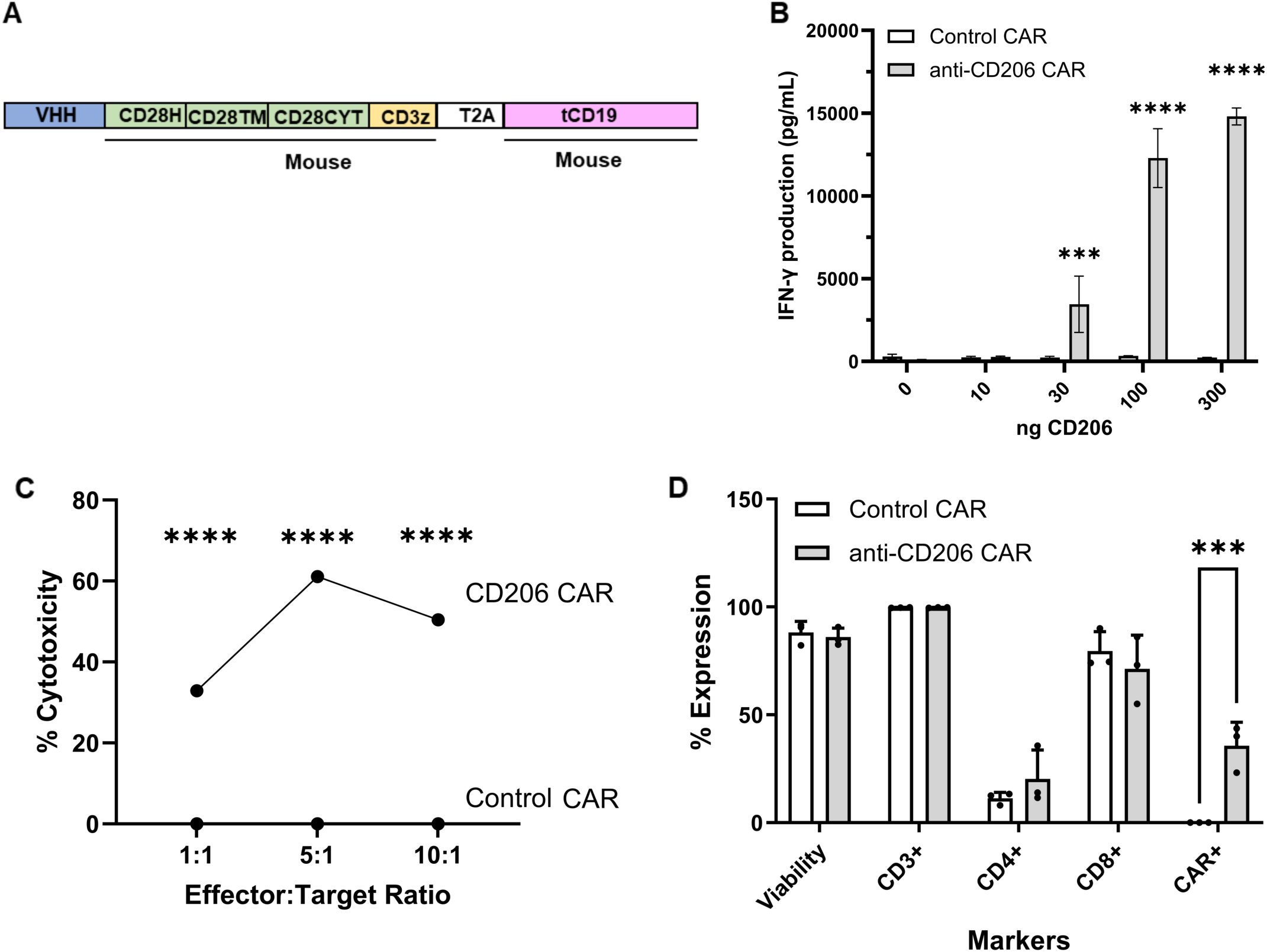
*In vitro* evaluation of anti-CD206 CAR T cells. (**A**) Construct design: Mouse CD28 is divided in hinge (H), transmembrane (TM) and cytoplasmic domains (CYT), followed by the CD3 zeta signaling domain. (**B**) Anti-CD206 or vector control-transduced CAR T cells were incubated with 0, 1, 3, 10, 30 and 100 ng/well recombinant CD206 protein for 24 hrs. Cell supernatants were collected and secreted IFN-γ was measured by ELISA (**C**) Cytotoxicity of transduced T cells (Effector) against mouse bone marrow cells (BM, Target). Levels of lactate dehydrogenase were measured after 6 hrs. (**D**) CD3, CD4, CD8, and CAR expression was measured in transduced cells at the end of process and prior to injection using flow cytometry. Error bars represent the SD of triplicates. *p≤0.05; **p≤0.01; ***p≤0.001; ****p≤0.0001.

We next evaluated anti-CD206 CAR T cells *in vivo*. Fibrosis was initiated on day 0 by delivery of bleomycin via osmotic minipump dorsal implantation. On day 6 post-bleomycin initiation, mice were either untreated or treated with vehicle (Hank’s balanced salt solution, HBSS), control CAR T cells, or anti-CD206 CAR T cells transferred by tail vein injection. Osmotic minipumps were removed on day 8, and skin was harvested at the conclusion of the experiment on day 21 (**Figure 3A**). We found that while anti-CD206 CAR T cell treatment did not significantly alter dermal thickness, DWAT was restored to control levels in mice that received anti-CD206 CAR T cells **(Figures 3B and C)**. Notably, administration of control CAR T cells failed to rescue DWAT loss.

**Figure 3.**
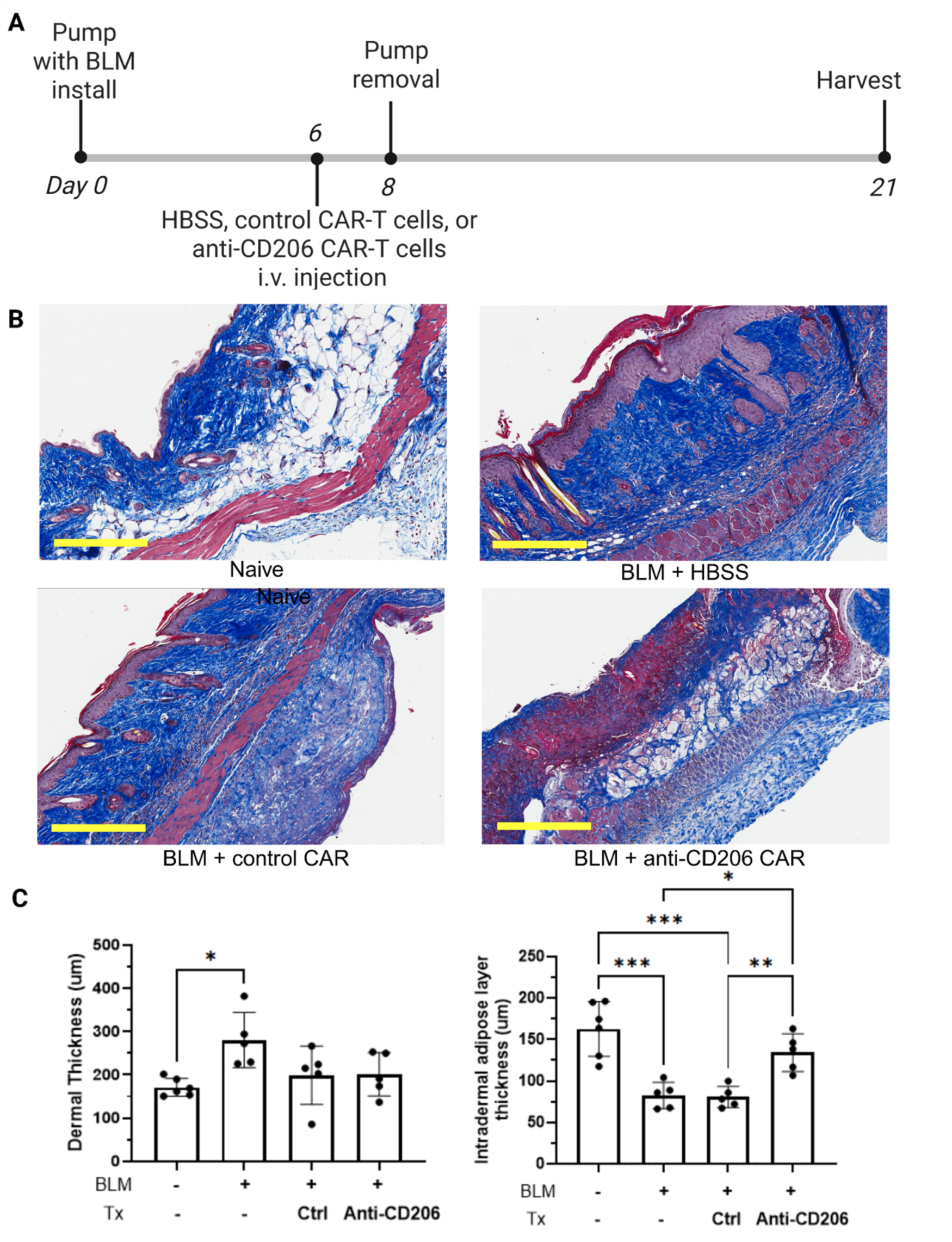
Systemic treatment with a single dose of anti-CD206 CAR T cells restores DWAT. (**A**) Experimental design: PBS or BLM treatment was initiated using osmotic minipumps on day 0 as above. Mice received a single dose of HBSS media, 3x10^6^ control CAR T cells or 3x10^6^ anti-CD206 CAR T cells via tail vein on day 6. Osmotic minipumps were removed at day 8. At the conclusion of the experiment at day 21, mice were euthanized and skin tissues were processed for histology. (**B**) Representative images of skin from control (PBS, non-BLM-treated) mice (top left), or fibrotic skin in mice administered BLM (top right), BLM and control CAR T cells (bottom left) or BLM and anti-CD206 CAR T cells (bottom right). Scale bar = 300μm. (**C**) Dermal (left) and adipose (right) thickness measurements were collected using Aperio Imagescope, where 5 measurements were collected per slide. *p≤0.05; **p≤0.01; ***p≤0.001; ****p≤0.0001.

### Anti-CD206 CAR T cell treatment remodels the dermal adipocyte and fibroblast landscape in bleomycin-treated mice

To interrogate how elimination of CD206^high^ macrophages alters the cellular heterogeneity of fibrotic skin, single nucleus RNA-seq (snRNA-seq) was performed on dermal tissues from mice treated with or without bleomycin and control or anti-CD206 CAR T cells as described in **Figure 3**. Initial analyses were conducted by integrating cells from all four treatment groups. Reference-based transfer learning leveraging two public datasets was used to generate putative assignments of dermal cell types, which were further refined using cluster-specific marker gene analysis coupled with expert review (see Methods). Importantly, all expected dermal cell types were recovered, including adipocytes.

Because fibroblasts are key mediators of SSc pathogenesis, subclustering was performed on the fibroblast population identified in **Figure 4A** to parse cellular heterogeneity. This analysis identified four fibroblast subclusters **(Figure 4B)**: myofibroblasts, denoted by expression of *Acta-2* (αSMA), lysyl oxidases (*Lox*, *Loxl2*, *Loxl3*), and ECM remodeling genes (*Col1a1*, *Tnc*, *Fn1*, and *Timp1*); hair follicle-associated fibroblasts, characterized by expression of *Col16a1*, *Mmp9*, *Twist1*, and *Apcdd1*; adventitial fibroblasts, which express *Pi16*, *Mfap5*, *Col14a1*, and *Cd248*; and papillary fibroblasts, marked by expression of *Wif1*, *Col6a5*, and *Mmp3*.

**Figure 4.**
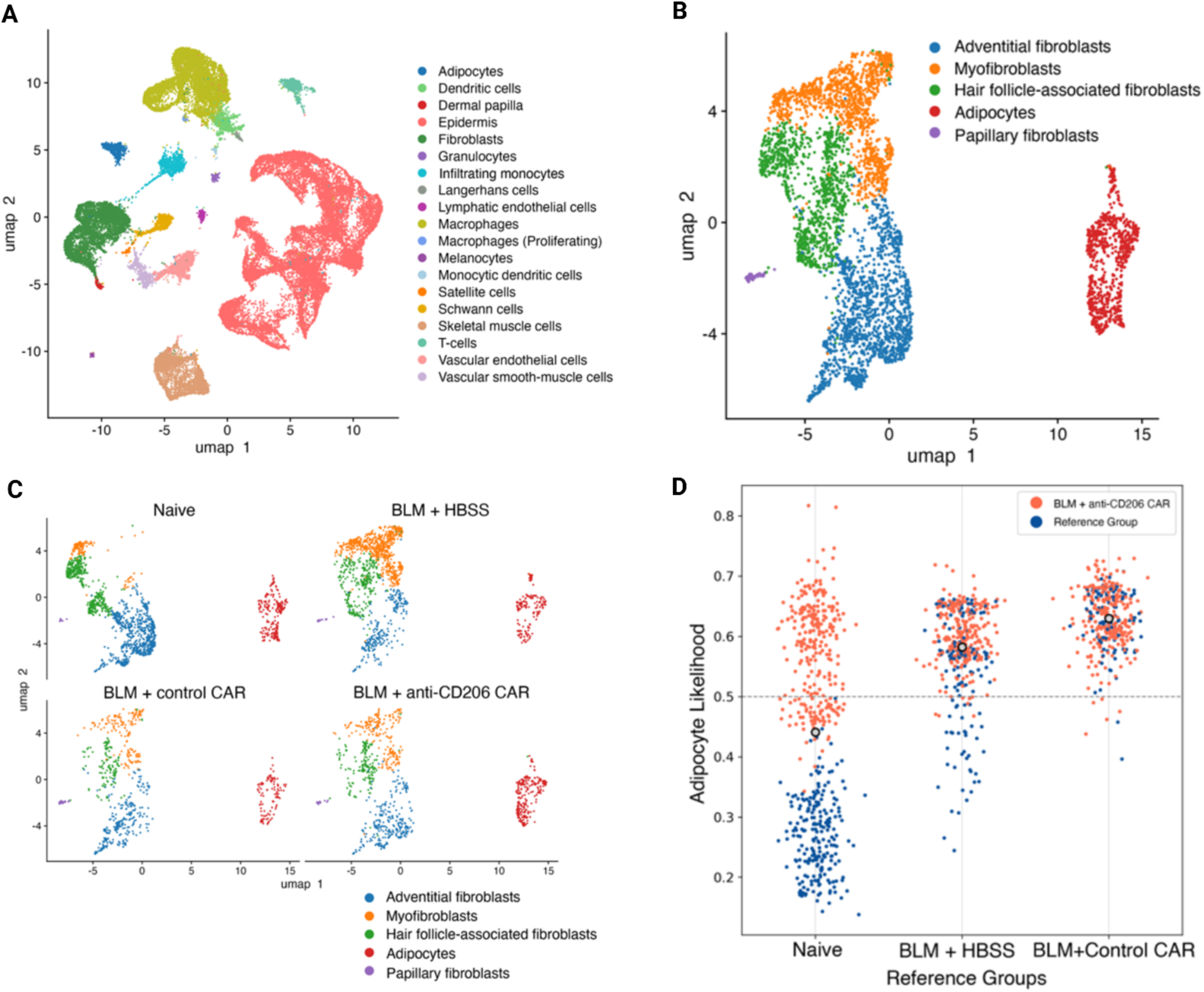
BLM and CAR T cell treatment modify fibroblast and adipocyte representation among dermal populations. A. UMAP representation of all dermal cell types identified, aggregated across all four treatment conditions. B. UMAP representation of fibroblast and adipocyte subclusters. C. Fibroblast and adipocyte subclusters on UMAP divided by treatment. D. anti-CD206 CAR T cell treatment-associated relative likelihood values from experimental perturbation analysis of all adipocyte cells. Relative likelihoods quantify effect of treatment with anti-CD206 CAR T cells vs. reference groups (Naïve, BLM+HBSS, or BLM+Control CAR). Cells with likelihood values >0.5 are more likely to be found in the BLM+anti-CD206 CAR T cell-treated sample, whereas cells with values <0.5 are more likely in the reference group. Color of points indicates sample of origin for each cell.

To gain greater insight as to how anti-CD206 CAR T cell immunotherapy modulates fibroblast subset representation in skin, each of these fibroblast subsets was analyzed for differences in cell type composition based on treatment condition **(Figure 4C)**. In accordance with expectations, bleomycin treatment was associated with expansion of myofibroblasts, which are central mediators of fibrosis and are largely absent from non-bleomycin treated skin. Control and anti-CD206 CAR T cell treatment resulted in attenuation of this population **(Figure 4C)**. Conversely, hair follicle-associated fibroblasts contracted in the presence of bleomycin treatment, consistent with follicular damage and alopecia that are observed with fibrosis [21]. Adventitial fibroblasts, which maintain the structural integrity and function of the vasculature, were similarly reduced with bleomycin administration. CAR T cell treatment did not restore these populations **(Figure 4C)**.

Given the observation that targeted elimination of CD206^high^ macrophages was associated with DWAT restoration, we next interrogated the effect of immunotherapy on dermal adipocyte representation. The mature dermal adipocyte population observed in non-bleomycin-treated mice, characterized by expression of adipokines *Adipoq* and *Retn* and regulators of fatty acid transport (FABP4), was reduced with bleomycin treatment, consistent with elimination of DWAT observed in **Figure 3**. However, administration of anti-CD206 CAR T cell therapy dramatically enhanced representation of the adipocyte cluster **(Figure 4C)**. Notably, control CAR T cell treatment failed to induce this change. To quantify differential abundance of cell types across treatment conditions, we used MELD (manifold enhancement of latent dimensions), which estimates the likelihood of cell types appearing in a specific treatment condition compared to a reference group. MELD was used to estimate the likelihood of cells being present in the skin sample derived from mice that received BLM+anti-CD206 CAR T cell treatment relative to each treatment group separately **(Supplementary Figure S5)**. When compared to treatment-naïve skin, relative likelihood values for adipocytes in mice that received BLM+anti-CD206 CAR T cells demonstrated a wide range of values, suggesting similar abundance of adipocytes in naïve and anti-CD206 CAR T cell-treated samples **(Figure 4D)**. However, when compared against skin from mice that received BLM+HBSS or BLM+Control CAR T cells, relative likelihood values of adipocytes in skin from BLM+anti-CD206 CAR T-treated mice were generally >0.5, suggesting enrichment of adipocytes in the BLM+anti-CD206 CAR T cell-treated sample **(Figure 4D, Supplementary Figure S5)**. Collectively, these data implicate a role for CD206^high^ macrophages in the regulation of adipocyte commitment and/or differentiation.

### CD206^+^ macrophages induce profibrotic skewing of adipose-derived stem cells (ADSCs)

We next interrogated the effect of CD206^high^ macrophages on adipogenesis in fibrotic skin. Prior work has shown that adipocytes and adipocyte progenitors play a role in the development and progression of dermal fibrosis [12, 22], and CD206^+^ macrophages have been shown to regulate adipocyte progenitor proliferation and differentiation in models of obesity [23, 24].

To elucidate the molecular mechanism by which CD206^high^ macrophages regulate adipocyte commitment in SSc skin, an *ex vivo* model system was developed to mimic the dermal fibrotic microenvironment. Dorsal skin sections from naïve or bleomycin-treated mice were cultured *in vitro* for 36 hours to generate conditioned media (CM). Because macrophage activation is derived from local micro-environmental signals, BMDMs were incubated with normal skin-conditioned media (NSCM) or fibrotic skin-conditioned media (FSCM) for 3 days **(Figure 5A)** to mimic the local dermal tissue milieux to which normal or fibrotic skin macrophages are exposed. After 3 days, media containing NSCM or FSCM were removed from BMDMs and replaced with fresh DMEM in the absence of CM for an additional 48 hours. NSCM and FSCM-activated macrophages were characterized by flow cytometry and ELISA. As demonstrated in **Figure 5B and Supplementary Figure S1,** FCSM-activated BMDMs were phenotypically consistent with human pro-fibrotic SSc macrophages [6], as evidenced by upregulated CD206 surface expression and increased CCL2 and IL-6 protein secretion.

**Figure 5.**
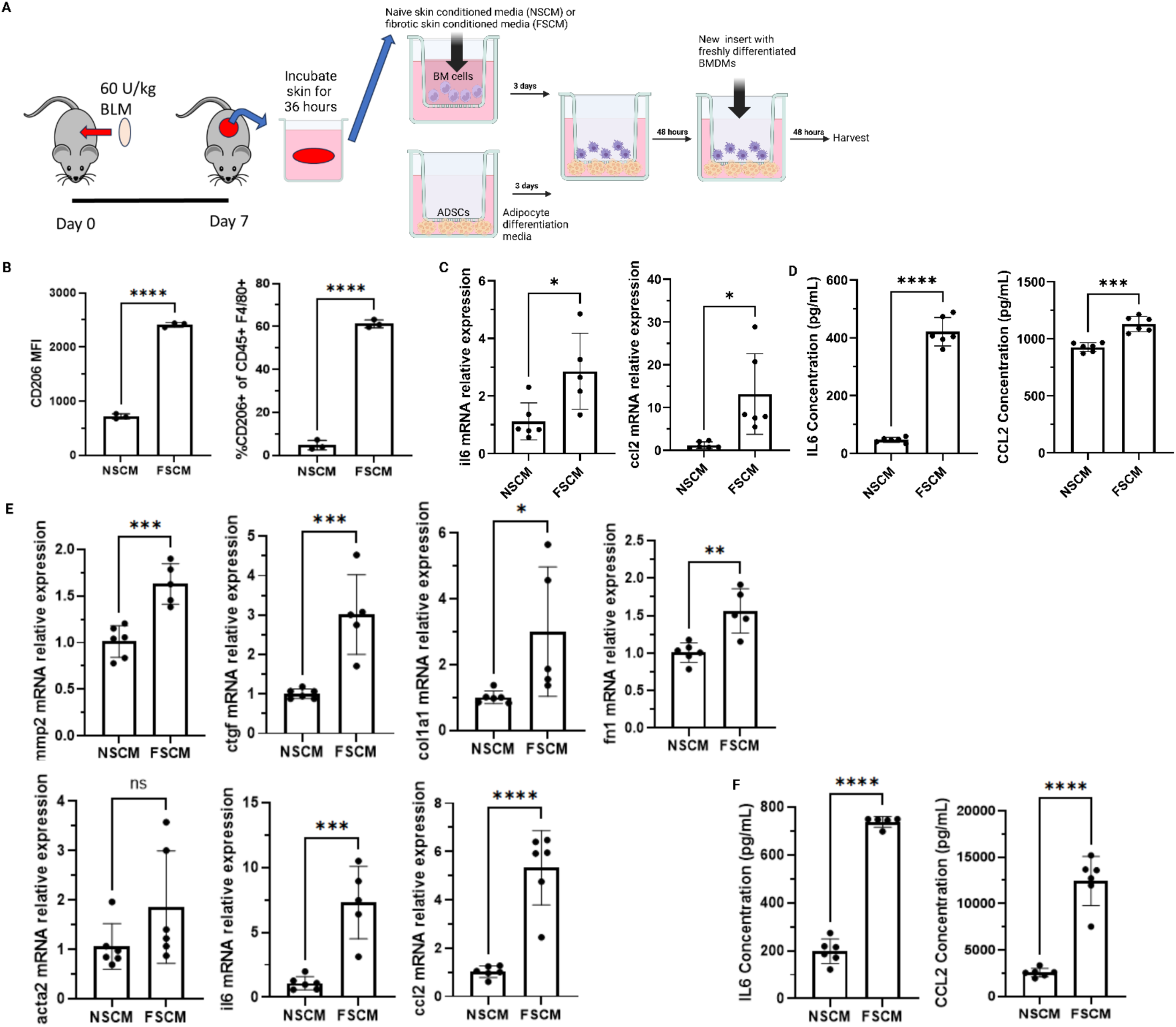
BMDMs activated with fibrotic skin-conditioned media (FSCM) induce fibrotic marker expression in ADSCs. **(A**) Schematic figure of experimental design. BMDMs were activated with CM from skin of normal mice (NSCM) or mice treated with BLM (FSCM) for three days. Cells were washed and media replaced with fresh complete DMEM media in the absence of CM and plated in upper chambers of Transwell plates. Primary ADSCs from C57B/L6 mice were isolated and plated in lower plate chambers and cultured in the absence of BMDMs for three days. CM-activated BMDMs and ADSCs were co-cultured for 48 or 96 hours as indicated in Methods. RNA from ADSCs was isolated and analyzed by qRT-PCR after 96 hours of co-culture. (**B**) BMDMs activated with NSCM or FSCM for three days were characterized (**B**) using flow cytometry to measure CD206 surface expression (**C**) using qRT-PCR to assess mRNA levels of IL-6 and CCL2. (**D**) IL-6 and CCL2 were quantified in CM-activated BMDM culture supernatants by ELISA. (**E**) Fibrotic gene expression was evaluated in ADSCs co-cultured with CM-activated BMDMs using qRT-PCR. (**F**) IL-6 and CCL2 levels were measured in co-culture supernatants using ELISA. *p≤0.05; **p≤0.01; ***p≤0.001; ****p≤0.0001.

To generate ADSCs, CD34^+^Sca1^+^CD24^+^ adipocyte progenitors were isolated from inguinal fat pads using flow cytometry **(Supplementary Figure S2A and B).** These selection criteria were based on prior work demonstrating that CD34^+^Sca1^+^CD24^+^ adipocyte progenitors can reconstitute the adipose layer in lipodystrophic mice [22]. We found that monoculture of ADSCs in differentiation media led to a significant increase in CD24 mean fluorescence intensity (MFI) in the CD34^+^Sca1^+^ ADSC population (Supplementary Figure S2C). Thus, the ADSC population used in the *ex vivo* culture system closely resembles the adipocyte progenitor population *in vivo*.

NSCM or FSCM-activated macrophages were then co-cultured with ADSCs in the absence of CM in wells separated with Transwell inserts for 48 hours **(Supplementary Figure S3)**. After this initial co-culture period, upper inserts with BMDMs were replaced with freshly-activated BMDMs and co-cultured with adipocyte precursors, again in the absence of CM, for an additional 48 hours **(Figure 5A)**. After a total of 96 hours of co-culture, RNA was isolated from ADSCs and BMDMs and analyzed for expression of profibrotic and inflammatory mediators. As demonstrated in **Figure 5C**, mRNA levels of matrix metallopeptidase 2 (MMP2), connective tissue growth factor (CTGF), fibronectin 1 (FN1), and type 1 collagen alpha 1 chain (COL1A1) increased significantly in ADSCs co-cultured with FSCM-differentiated BMDMs. Notably, these are hallmark genes characteristic of early stage fibroblasts [24, 25], while expression of α-SMA, which is elevated in mature myofibroblasts [26], was not significantly altered. Consistent with increased expression of ECM mediators, IL-6 and CCL2 mRNA and protein expression were also significantly increased in co-cultured ADSCs **(Figures 5C and 5D).** Collectively, these results implicate a role for CD206^+^ macrophages in the redirection of ADSC commitment from adipocyte to fibroblast.

### CD206^+^ macrophage-derived IL-6 regulates ADSC fibrotic commitment

IL-6 has been implicated as a regulator of stemness and de-differentiation [27, 28], is overexpressed by SSc macrophages [6, 7], and is a well-characterized inducer of fibrotic activation [29]. Because IL-6 production was significantly increased in FSCM-activated BMDM and ADSC co-cultures, we interrogated a potential role for IL-6 in ADSC lineage commitment. Co-cultures were established as in **Figure 6A** in the presence or absence of blocking anti-IL-6 receptor antibody, and markers of adipocytic or fibrotic commitment were evaluated. IL-6 receptor blockade significantly increased levels of ADIPOQ, which encodes adiponectin and is expressed by mature adipocytes [30], but decreased expression of fibroblast-associated FN1 [31] **(Figure 6B and 6C).** In contrast, blockade of TGF-β failed to elicit changes in ADSC commitment **(Supplementary Figure S4)**. Collectively, these results suggest that CD206^+^ macrophage-derived IL-6 contributes, at least in part, to driving ADSC fibroblast-like commitment in the context of fibrotic disease.

**Figure 6.**
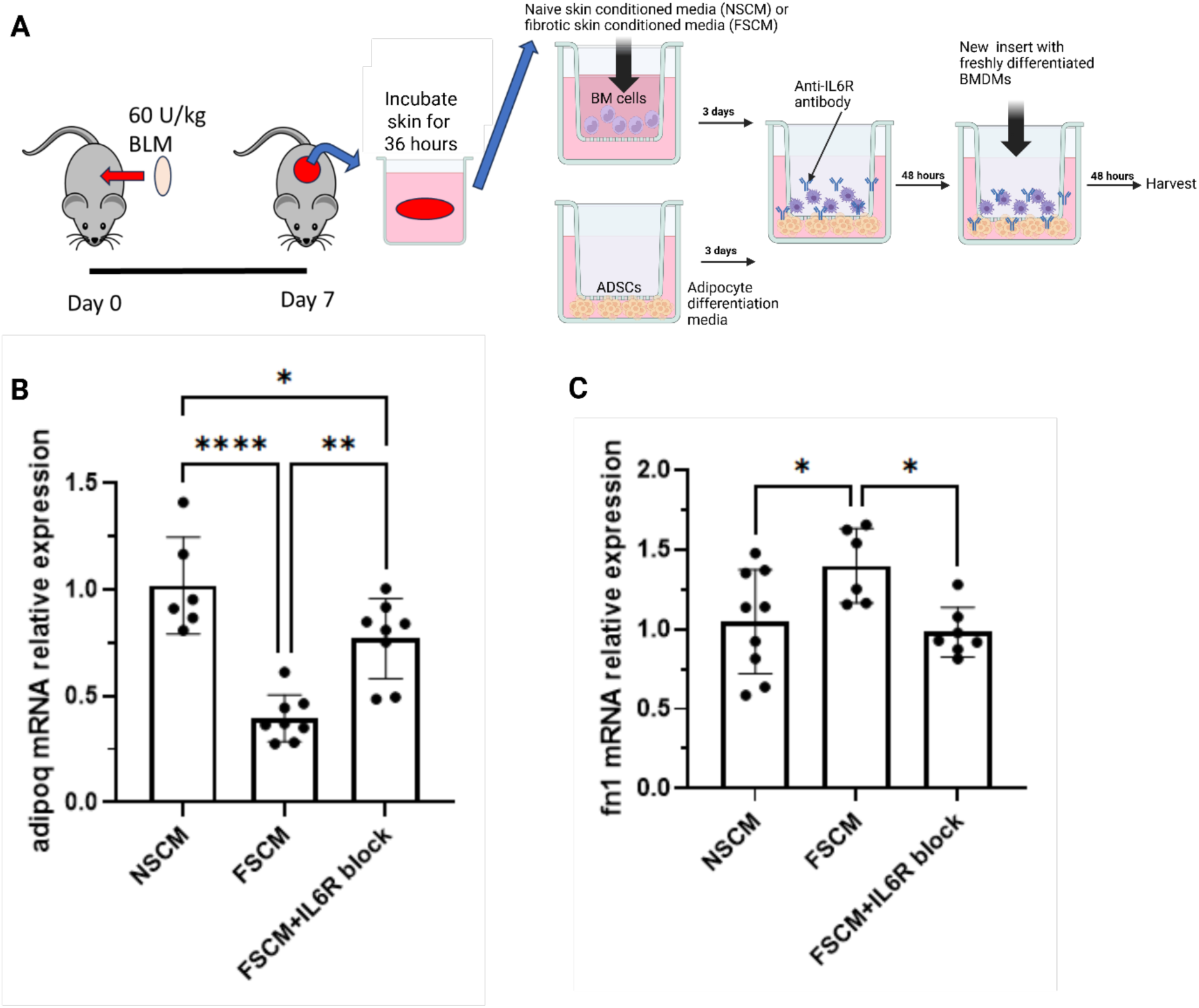
IL-6R blockade redirects ADSC lineage commitment. (**A**) Experimental design: Co-cultures were established as in 5A in the presence or absence of IL-6R blockade. 100 μg/mL anti-mouse IL-6R monoclonal antibody was added co-cultures for 96 hrs. RNA was isolated from ADSCs and expression of (**B**) adipocytic and (**C**) fibrotic genes was measured using qRT-PCR. *p≤0.05; **p≤0.01; ***p≤0.001; ****p≤0.0001.

## Discussion

Because prior work by our group and others identified a central role for CD206^high^ macrophages in the pathogenesis of SSc, we developed a cellular immunotherapeutic approach to eliminate these cells. While B cell-targeting (anti-CD19) CAR T cells have shown significant therapeutic promise in the treatment of autoimmune diseases [32-34], our report is the first to use CAR T cells to attack SSc profibrotic macrophages, a prime driver of SSc pathogenesis. We now demonstrate the potential therapeutic efficacy of using CAR T cells to target dermal macrophages and have identified a novel role for CD206^high^ macrophages in regulation of SSc skin pathogenesis.

Loss of DWAT is a consistent hallmark of SSc that occurs early in disease, preceding the development of fibrosis [12]. Prior work in the bleomycin model implicates altered adipocyte to fibroblast cell fate transition driven by TGF-β as an underlying regulator of DWAT loss in SSc [12], although the molecular mechanism by which this occurs is incompletely understood. We now demonstrate that targeted elimination of CD206^high^ macrophages restores DWAT, suggesting that CD206^high^ macrophages provide key signals that shift adipocyte-fibroblast cell lineage commitment. Moreover, we demonstrate that ADSCs co-cultured with CD206^high^ macrophages are characterized by increased expression of fibrogenic genes and suppression of adipogenic genes. This is driven, at least in part, by IL-6, as blockade with anti-IL-6 receptor antibody results in restoration of ADIPOQ gene expression to levels not significantly different from ADSCs co-cultured with BMDMs activated with NSCM. A significant shift in immune activation also developed with anti-CD206 immunotherapy, with decreased inflammatory monocytic dermal infiltration and enhanced representation of a myeloid population associated with relief of oxidative stress. Furthermore, myofibroblasts were reduced by anti-CD206 CAR T cell treatment, suggesting CD206^high^ macrophage elimination remodels the SSc dermal micro-environment.

In addition to DWAT loss, SSc is also characterized by reductions in circulating levels of adiponectin, an adipokine that regulates metabolism, inflammation, and fibrosis [35, 36]. Adiponectin signaling is diminished in lesional skin biopsies derived from individuals with SSc and is negatively correlated with disease severity and activity. Moreover, overexpression of adiponectin in bleomycin-treated mice is associated with significant attenuation of fibrosis [37], suggesting a causal link between adipocyte loss and fibrosis. Thus, we hypothesize that CD206^high^ macrophage elimination results not only in attenuation of DWAT loss but also diminished fibrosis as adipocytes are replenished and adiponectin production is restored. In our model, CD206^high^ macrophages provide signals mediated by IL-6 that alter ADSC commitment from an adipogenic to a fibrogenic cell lineage; activated fibroblasts then secrete ECM and inflammatory mediators, including IL-6, culminating in fibrotic disease. Because of adipocyte loss, adiponectin levels are reduced and the normal regulatory control mechanisms mediated by this adipokine to inhibit fibrosis are thwarted, resulting in sustained fibrotic activation and decreased adipocyte commitment. Elimination of CD206^high^ macrophages reduces DWAT and adiponectin loss, leading to both acute and sustained attenuation of fibrosis. Additional experiments are underway to assess long-term effects on DWAT restoration and fibrosis. We are also assessing the effect of anti-CD206 CAR T cell treatment on other end-target organs affected by fibrosis in SSc, including lungs and heart, as CD206^high^ macrophages have been implicated as mediators of fibrosis at these sites [20, 38].

While ample evidence supports elevated CD206 expression on SSc macrophages and in fibrotic disease [6, 7, 20, 39, 40], CD206 is also expressed at low levels on normal myeloid cells and discrete populations of liver endothelial cells. Since CAR T cells are highly sensitive to antigen, on-target effects on normal (non-diseased) tissue are an important consideration. Notably, we have not discerned adverse events in these studies. The peak observed CAR T cell cytotoxicity against CD206-expressing MØs was ∼60% **(Figure 2C)**, implying that killing was restricted to CD206^high^ cells, which may explain why we failed to observe effects on non-pathological tissue. Nonetheless, it will be imperative to monitor cytokine release and potential off-target or on-target-off-tissue effects as experimental parameters are altered.

Attesting to the importance of CD206^+^ macrophages in fibrotic disease, additional therapeutics aimed at modulating macrophage activation have indicated potential therapeutic efficacy in both mouse models of fibrosis and in human cell culture. For example, nintedanib and pirfenidone inhibit expression of surface CD206 on murine and human monocyte-derived macrophages [41, 42], and the use of a synthetic peptide that binds CD206, RP-832c, has been shown to inhibit pulmonary fibrosis in bleomycin-treated mice [43]. However, pharmacological and peptide inhibitors may be associated with significant adverse effects and may fail to specifically target cellular mediators. In contrast, cellular therapy allows for a more personalized approach by using each patient’s own T cells to generate CAR T cells and allows for incorporation of additional therapeutics to be delivered to the diseased tissues more specifically. This makes the CAR T cell approach to targeting CD206^high^ macrophages potentially more effective and less likely to be associated with the side effects often observed with non-cellular therapeutic treatments.

To assess potential therapeutic efficacy, this study was designed to test the effect of a single dose of anti-CD206 CAR T cells on established fibrosis. Accordingly, CAR T cells were administered systemically one-week post-initiation of bleomycin treatment. However, it is possible that enhanced effects on fibrotic inhibition would be observed with repeated administration and/or by altering CAR T cell dose. Optimization of anti-CD206 CAR T cell treatment is a primary focus of our current studies. Nonetheless, the demonstration that systemic delivery of a single dose of anti-CD206 CAR T cells results in significant disease amelioration suggests this approach may be therapeutically beneficial by attacking fibrosis at a central node of pathogenesis.

Given SSc heterogeneity, it is likely that different patient subsets will respond differentially to anti-CD206 CAR T cell treatment. In this regard, prior work has shown substantial heterogeneity in SSc patient response to immunomodulatory therapies like mycophenolate mofetil (MMF) [44]. Furthermore, the bleomycin mouse model may not represent the full spectrum of SSc patients [45]. Accordingly, we plan to perform additional studies in additional mouse models of disease, including the sclGVHD model. Future studies may include work with humanized mouse models of disease [46]. One caveat of this study is evaluation of CAR T cell persistence in skin due to limitations associated with detection. To address this, we plan to incorporate a GFP tag into anti-CD206 CAR T cell constructs that will permit live imaging during treatment. Additionally, the effect of CAR T cell therapy on other SSc-affected tissues is unknown. Nevertheless, this work establishes, for the first time, the therapeutic feasibility and efficacy of using CAR T cell immunotherapy to target macrophages, a key mediator of fibrotic activation, in the treatment of SSc skin disease.

Collectively, these results identify a novel role for CD206^high^ macrophages in SSc pathogenesis and suggest that targeting CD206^high^ macrophages with CAR T cells may provide significant therapeutic benefit in not only delaying the progression of fibrosis, but also in the restoration of fibrotic skin to a normal, non-diseased state. Because the mechanisms by which macrophages regulate fibrosis in SSs are common to other fibrotic diseases, this approach may have significant therapeutic benefit in the treatment of not only SSc but also in other conditions characterized by pathological CD206^high^ macrophages.

## Materials and Methods

### Sex as a biological variable

Because SSc disproportionately affects women, with estimates of female to make ratios ranging from 3:1-10:1 [47], female mice were used in all experiments included in this study.

### Mice

Mice were handled ethically according to the Regulations for the Management of Laboratory Animals at the Geisel School of Medicine at Dartmouth. The experimental protocol for the ethical use of these animals was approved by the Institutional Animal Care and Use Committee at the Geisel School of Medicine at Dartmouth (protocol 00002183). 6-7-week-old C57BL/6 female mice (Jackson Laboratory) were housed for a week prior to use in experiments. Phosphate buffered saline (PBS) or bleomycin was administered to mice using osmotic minipumps (Azlet) that delivered bleomycin at a rate of 0.5 μl/hr for 7 days as described [48]. Bleomycin was dissolved in PBS at a concentration of 60U/kg. On day 0, surgical sites were shaved and minipumps were implanted by making small incisions under the back skin of mice anesthetized with isoflurane. Incisions were closed using surgical staples. Mice were treated with carprofen at the time of and 24 hours post-surgery for pain management. Mice were monitored and weighed for the entire experimental duration. Pumps were removed 8 days post-implantation.

For experiments in **Figure 1**, 200μl of 5mg/ml clodronate or empty control liposomes (Liposoma) were administered via intraperitoneal injection starting 3 days before osmotic pump insertion and continued every 3 days thereafter until the conclusion of the experiment as outlined in **Figure 1D**. Body weight and food intake were monitored daily. At the conclusion of experiments (day 21), mice were euthanized and dermal tissues harvested for measurements of dermal and adipose layer thickness and histopathological analysis.

To evaluate the effect of targeted elimination of CD206^high^ macrophages, control or anti-CD206 CAR T cells (3x10^6^) were resuspended in HBSS and administered intravenously via tail vein injection using 27-gauge needle on day 8. Mice were treated with HBSS alone in the presence of bleomycin as negative control. Dermal tissues were harvested from mice at day 21 for analysis of dermal and DWAT thickness and histopathology (H&E staining); a set of tissues was formalin-fixed for sn-seq analysis.

### Retroviral packaging and transfection

HEK 293T cells were cultured in Dulbecco’s modified Eagle (DMEM) (Cytiva SH30243.02) media supplemented with 10% heat-inactive FBS (Cytiva SH30071.02), passaged one to two times at <60% confluency in the absence of Pen-Strep, and then seeded onto 10-cm culture dishes at a density of 2.0 x 10^6^ cells per dish and cultured for ∼36 hours to 80-90% confluency. One hour before transfection, cells were provided with fresh media. DNA transfections were performed using polyethylenimine (PEI). Briefly, 5 mg of CAR or cytokine plasmid were combined with 5 mg of pCL-Eco packaging plasmid (Addgene 12371) in 600 μL of OptiMEM (Gibco 31985062). To this mixture, 30 μL of PEI (1mg/mL) was added and incubated for 10 minutes at room temperature prior to dropwise addition to HEK 293T cells. The media was replaced with 5 ml with fresh media including Pen-Strep 18 hours post-transfection. Viral supernatants were harvested at 48- and 72-hours post-transfection, filtered through a 0.45 mm filter and used immediately or stored at -80°C.

### CAR T cell generation

Splenocytes from CD45.1^+^ B6 mice were activated using Concanavalin A (2.5 mg/mL; Sigma C5275) in cRPMI media (RPMI-1640 (Cytiva SH30255.01), 10% heat-inactive FBS, 10 mmol/L HEPES (Gibco 15630080), 100 mmol/L nonessential amino acids (Gibco 11140050), 1 mmol/L sodium pyruvate (Gibco 11360070), 100 U/mL penicillin (Gibco 15140122), 100 mg/mL streptomycin (Gibco 15140122), and 50 mmol/L of 2-mercaptoethanol (Gibco 21985023). Following 24-48 hours, activated T cells were retrovirally transduced to express anti-CD206 CAR by seeding 8.0 x 10^6^ cells per well in a 24-well plate, resuspension in viral supernatant and polybrene (1 mg/mL; Sigma 107689), followed by spinoculation at 1,500xg for 90 minutes at 37°C. The viral supernatant was subsequently removed, and cells were maintained in culture at a concentration of 10^6^ cells/mL in cRPMI with 25 U/mL recombinant human IL-2 (NCI-Frederick BRB Repository). Flow cytometric analysis was conducted 72 hours post-transduction, and CAR expression levels were evaluated prior to CAR T cell adoptive transfer.

### snRNA-seq and bioinformatic analysis

snRNA-seq analysis of Formalin Fixed Paraffin Embedded (FFPE) skin tissues was performed using the 10x Genomics Flex method following Demonstrated Protocol CG000784 and User Guide CG000527. Briefly, 2 x 25uM sections were cut from each FFPE block into a 1.5mL tube and stored at 4C for no more than 24hrs prior to processing. Sections were transferred to GentleMACS C Tubes (Miltenyi), deparaffinized in xylene, rehydrated in graded alcohols and dissociated in 1mg/ml Liberase TM enzyme using the recommended GentleMACS program. Dissociated tissues were passed through 30um filters, and nuclei were counted using AO/PI fluorescence on a Luna FX7 automated cell counter. Nuclei were hybridized to whole-transcriptome probe sets and loaded onto a Chromium X instrument, targeting 10,000 nuclei for capture per sample. The resulting gene expression libraries were sequenced on a NextSeq2000 (Illumina) targeting 25,000 reads/cell.

### snRNA-seq preprocessing

Raw sequencing data in FASTQ format were processed using the Cell Ranger (v.8.0.0) *multi* pipeline to construct a cell-by-gene matrix of raw counts, using probe-set v.1.0.1 and reference genome (mm10-2020-A). R-package Seurat (v.5.0.0) [49, 50] was used for quality control, normalization, and integration. Data was imported into R (v.4.4.1) using Seurat function *Read10X* with arguments min.cells = 10, min.features = 50. Droplets were then filtered to retain those with ≥1000 UMIs, ≥300 genes, ≤10% reads aligned to mitochondrial genes. Datasets from each sample were normalized separately using function *SCTransform* [51] with default arguments and then integrated to remove batch effects using the Seurat integration workflow [52] with reciprocal PCA to identify anchors. Genes to use during integration were selected from each dataset using function *SelectIntegrationFeatures* with argument n_features = 200. Datasets were then processed using function *PrepSCTIntegration* and subject to PCA analysis with function runPCA. Integration anchors between the datasets were identified using *FindIntegrationAnchors* with arguments normalization.method = “SCT”, dims = 1:50, reduction = “rpca”. Datasets were then integrated with the *IntegrateData* function, again with normalization.method = “SCT”. Integrated data were projected into the same low dimensional space using functions *RunPCA* and *RunUMAP* using the top 40 principal components. Function *FindClusters* was used to construct a shared nearest neighbor graph, which was then used to identify unsupervised clusters across several resolutions using function FindClusters with the Louvain algorithm. R-package clustree [53] was used to examine cluster stability across all clustering resolutions, leading to selection of 0.3 as a preliminary resolution with which to begin cell type annotation.

### Cell-type annotation

Cell types were annotated using a combination of reference-based transfer learning and cluster-specific marker gene review. For reference-based annotation, two publicly available reference datasets were used: a skin-specific atlas of cell types present during hair growth and rest [54] (GSE129218) and PanSci [55] (GSE247719), a large multi-tissue atlas comprising 21,786,931 cells. The motivation for use of the PanSci reference was to provide additional support for annotations derived from the smaller skin-specific atlas (5,767 cells) and facilitate annotation of additional stromal and immune cell types not observed in the skin-specific atlas. For the skin-specific reference, Market Exchange (MEX) format files were downloaded from GSE129218 and read into R using function *Read10X*. Data from the PanSci reference was downloaded in H5ad format from GSE247719 and read into python (v.3.11.8) with scanpy (v.1.11.1) function *read_h5ad* for preprocessing. Data were filtered to include cells from only 3-month-old mice and wild-type genotypes. Due to the size of the PanSci reference, the data was subset to 50,000 cells that effectively summarize the heterogeneity of transcriptomic profiles in the full dataset using geometric sketching [56]. The sketching procedure was performed separately on cells from each tissue, with the number of cells selected per tissue being directedly proportional to the fraction of the input dataset. Tissue-specific datasets were combined with function *concat* and written to h5 files with function write_h5ad, before reading into R using function *LoadH5Seurat*. Raw read counts for both reference datasets were normalized using *SCTransform* [51] and integrated across samples to remove batch effects, using the same procedure described above. Anchors between the reference and query datasets were identified using function *FindTransferAnchors* with the first 30 dimensions and normalization.method = “SCT”. Labels were transferred to the query dataset using function *TransferData*. Prediction scores were reviewed for each transferred cell type to assess confidence and specificity of transferred labels. Cells with prediction scores of less than 0.5 were labelled as “Unknown” and removed from the dataset. Dimensionality reduction and unsupervised clustering was recomputed using the remaining cells. Unsupervised clusters determined at a resolution of 0.3 (as described above) were then assigned to cell types using a ‘majority rule’ approach [57], where clusters labels were assigned based on the cell type indicated by the majority of cells in each cluster. In cases where the reference-based annotations suggested subclusters were present, unsupervised clusters from deeper resolutions were reviewed to identify clusters composed predominantly of a single cell type. Marker genes were used to confirm reference-based annotations, resolve clusters composed of mixed cell types, or annotate ambiguous clusters. To detect marker genes, counts from the ‘RNA’ assay of the Seurat object were normalized with function NormalizeData and provided to function *FindMarkers* with arguments min.pct = 0.25 and only.pos = TRUE. Only markers with log2 fold-change values >1 and Bonferroni-adjusted *P*-values <0.05 were considered.

### Experimental perturbation analysis

We used the MELD algorithm (v.1.0.2) [58] to assess the effect of experimental perturbations (i.e. bleomycin or CAR T cell treatment) on the abundance of annotated cell types. In contrast to traditional differential abundance methods, MELD estimates a manifold from all cells across experimental conditions and uses this to estimate a likelihood that each cell would be observed in either the treatment or control condition. We used cells from the bleomycin and anti-CD206 CAR T cell-treated sample (BLM+anti-CD206 CAR) as the ‘treatment’ group and compared these to cells from ‘control’ samples (Naïve, BLM+HBSS, or BLM+Control CAR). Cells with a likelihood of >0.5 were more likely to be found in the anti-CD206 CAR T cell-treated sample, whereas cells with a likelihood of <0.5 are more likely to be found in the respective control sample. Examination of these ratios grouped by annotated cell type allows us to identify cell types more likely to be present under specific experimental conditions.

### Sub-clustering analysis

To perform sub-clustering analysis of populations of interest, raw data were read into R as described above and filtered to specific cell types labeled in the annotation of the complete dataset. These data were then subject to normalization, integration, dimensionality reduction, clustering, and marker detection as described above. Upon initial review of cluster-specific marker genes, a small population of contaminating keratinocytes was identified and removed from downstream analysis. A clustering resolution of 0.4 was ultimately selected for downstream analysis, and custom marker gene review was used to assign clusters to specific cell types.

### BMDM and ADSC isolation and culture

Bone marrow cells were flushed from femurs and tibias of 8-week-old C57BL/6 female mice (Jackson Laboratory) using syringes fitted with 18-gauge needles in cold RPMI. Cells were passed through 70 μm filters and washed in cold DMEM. BMDMs were subsequently cultured in complete DMEM containing 10% FBS and penicillin/streptomycin (pen/strep) and 20 ng/ml M-CSF as described [59].

ADSCs were obtained from inguinal fat pads of 8-week-old C57BL/6 female mice. Tissues were mechanically minced and digested with collagenase containing BSA fraction V, collagenase D, dispase II, and calcium chloride in PBS for 30 minutes at 37C. Following digestion, cells were resuspended after centrifugation at 1500 rpm and cell pellets were washed with fresh DMEM and 10% FBS to remove residual enzyme post-digestion. Cells were analyzed using flow cytometry and ADSCs were sorted based on CD34/Sca-1 dual positivity as outlined in **Supplementary Figure S2**.

### Co-culture experiments

Bone marrow cells were plated in the upper chambers of Transwell plates in DMEM with 50% normal (NSCM) or fibrotic (FSCM) skin conditioned media. Dermal conditioned media (CM) was generated by excising the fibrotic skin from mice with implanted with osmotic pumps containing saline or bleomycin followed by incubation in DMEM for 36 hours. ASDCs were plated in the lower chambers of Transwell plates in differentiation media (DMEM/F12 1:1 media with IBMX, dexamethasone, insulin, rosiglitazone, FBS and pen/strep). For antibody blockade experiments, either 100 mg/mL anti-mouse IL-6R monoclonal antibody (BioXCell) or 2 mg/ml anti-TGF-β pan-neutralizing antibody 1D11 (Invitrogen) was added to Transwell plates for the duration of co-culture.

### RNA isolation, cDNA synthesis, and qRT-PCR

Total RNA was isolated from ADSCs using the Quick-RNA Microprep Kit (Zymo, Cat. #11-328M) per the manufacturer’s instructions. Complementary DNA was synthesized using qScript™ Ultra cDNA SuperMix (QuantaBio, Cat. #955161–100). Quantitative real time PCR (qRT-PCR) was performed using TaqMan Probes single-tube assays (Life Technologies, Cat. #4324018) for murine MMP9, CTGF, COL1A1, FN1, ACTA2, IL6, CCL2, and ADOPIQ. The StepOnePlus Real-Time PCR System (Applied Biosystems) was used for amplification and detection. mRNA levels were normalized to ACTB, and thermal cycling conditions were performed as described [6].

### Flow cytometry

Plates were spun at 300 x g for 5 minutes to pellet cells. Cells were placed on ice, washed 1x with PBS and stained with LIVE/DEAD™ Fixable Yellow (Invitrogen, Cat.#L34959) for 15 minutes on ice prior to blockade with mouse Fc blocking antibodies. Surface markers were stained for 20min on ice in flow buffer (1X PBS, 2% BSA, 2mM EDTA). Doublets were excluded using forward scatter A (FSC-A) vs. FSC-H gating. Cells were analyzed using a MACSQuant 8-color Cytometer (Miltenyi) with 5 laser sources and FlowJo 10.8.1 software.

### ELISA

Cell-free supernatants were collected from cultures and secreted levels of murine CCL2 and IL-6 were measured by ELISA (Thermofisher). ELISA standard curves and calculations were performed using the Gain-Data tool by Arigo Biolaboratories.

### Histology

Tissues were fixed in 10% neutral buffered formalin, paraffin-embedded, and sliced parallel to the cassette to create a 4µm cross section of the center of the tissue prior to staining with hematoxylin and eosin (H&E). Sections were dried at room temperature before loading on the Tissue-Tek Prisma Stainer (Sakura Finetek USA) and running the automated routine H&E program. Tissue thickness was calculated using the average of five individual measurements at roughly equal intervals across the length of the tissue using the measure tool in Aperio ImageScope software.

### Statistical analysis

Results are described as mean (s.e.) and were analyzed by Welch’s t-test, one-way ANOVA or two-way ANOVA for multiple comparisons using GraphPad Prism 11. Significance was achieved at p < 0.05.

### Study approval

All experiments were conducted with the approval of the Institutional Animal Care and Use Committee at the Geisel School of Medicine at Dartmouth (protocol 00002183), and in compliance with the NIH Guidelines for Use and Care of Laboratory Animals.

## Data Availability

The raw data files for murine control and bleomycin-treated snRNA-seq samples will be deposited into the Gene Expression Omnibus (GEO) and made publicly available. All data generated during this study are included in this manuscript (and its supplementary information files). Additional detailed information is available from the corresponding author on request.

## Author Contributions

CP, MLW, and PAP designed the research. CP, AOM, RB, and JG performed and analyzed experiments. HJ, NNK, OW, and FWK performed snRNA-seq studies and analyses. YHH and JM provided expertise with CAR T cell studies (design, production and in vitro validation). All authors contributed to writing and editing of the manuscript. PAP provided overall guidance for the project.

## Funding

This work was supported by funding from the National Scleroderma Foundation (PAP), the Scleroderma Research Foundation (MLW), and NIH (R21AI169420 and R21AI178651, PAP and MLW; and P20 GM130454, MLW). Funding support was also received from Department of Defense grant W81XWH-21-1-0878 (MLW). CP was a recipient of the National Scleroderma Foundation Predoctoral Research Fellowship. This work is the result of NIH funding, in whole or in part, and is subject to the NIH Public Access Policy. Through acceptance of this federal funding, the NIH has been given a right to make the work publicly available in PubMed Central.

## Acknowledgements

We would like to acknowledge assistance from D. Tyler Boone with CAR T cell generation, and DartLab, the Immune Monitoring and Flow Cytometry Resource at the Dartmouth Cancer Cancer Center (NCI Cancer Center Support Grant 5P30CA023108-37) with flow cytometry studies. Research reported in this publication was supported through Geisel School of Medicine at Dartmouth’s Center for Quantitative Biology through a grant from the National Institute of General Medical Sciences of the National Institutes of Health under Award Number P20GM130454. Graphics were created using Biorender.

## Notes

Conflicts of Interest: Joana Murad-Mabaera, Michael L. Whitfield, and Patricia A. Pioli have submitted a patent application under 35 U.S.C. 371 based on International Application No. PCT/US2018/047101 for the use of anti-CD206 CAR T cell immunotherapy for treatment of fibrotic disease. MLW is a scientific founder of Celdara Medical LLC.

### Competing Interest Statement

Joana Murad-Mabaera, Michael L. Whitfield, and Patricia A. Pioli have submitted a patent application under 35 U.S.C. 371 based on International Application No. PCT/US2018/047101 for the use of anti-CD206 CAR T cell immunotherapy for treatment of fibrotic disease. MLW is a scientific founder of Celdara Medical LLC.

