## Supplementary material for "Anti-CD206 CAR T Cell Treatment Restores Fibrosis-Induced Loss of Dermal White Adipose Tissue": figures

**Supplementary Figures**

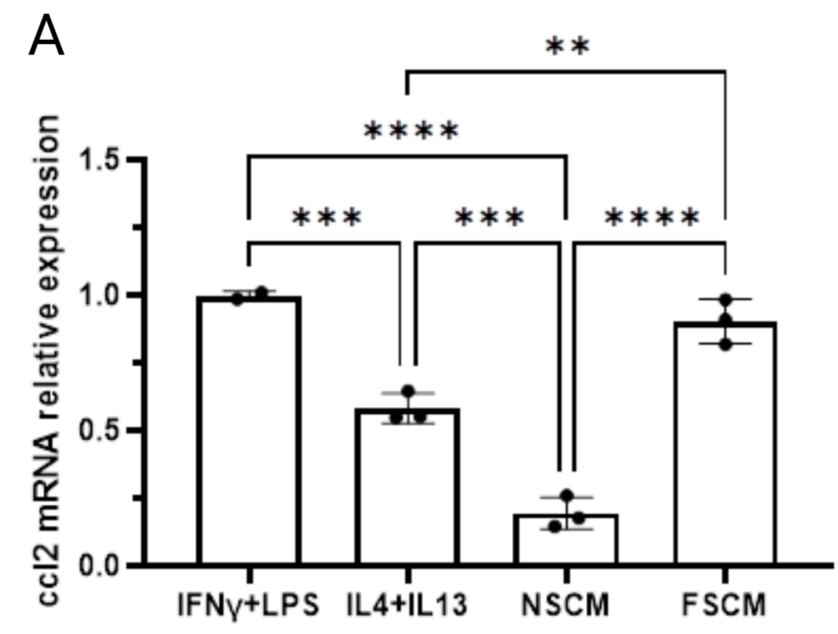


**Figure S1. Characterization of differentiated BMDM.** BMDMs were activated with IFN-γ/LPS; IL-4/IL-13; normal (non-BLM-treated) skin conditioned media (NSCM); or fibrotic skin conditioned media (FSCM) for three days and mRNA was extracted from the resulting populations. CCL2 mRNA expression was analyzed using qRT-PCR. *p≤0.05; **p≤0.01; ***p≤0.001; ****p≤0.0001

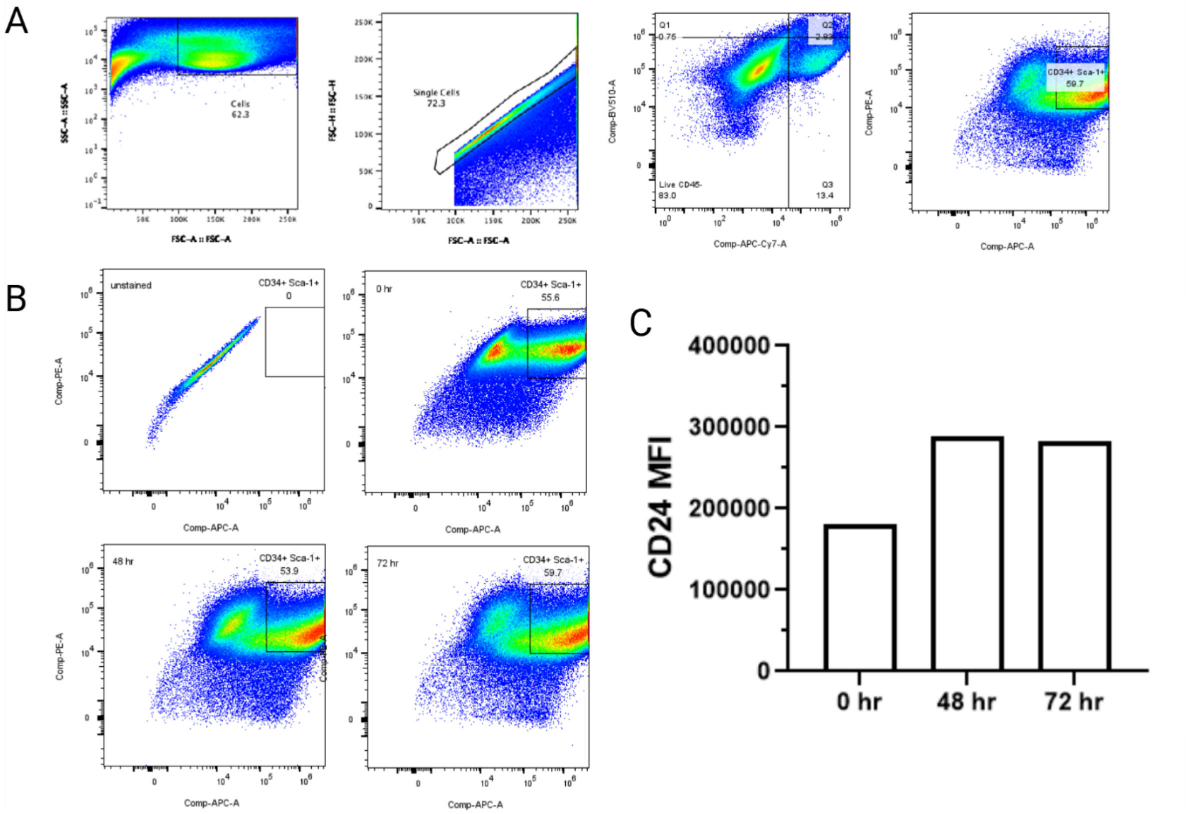

**Figure S2. Characterization of differentiated ADSCs.** ADSCs cultured for 0, 48, and 72 hrs in differentiation media were analyzed using flow cytometry. (**A**) Gating strategy for ADSCs. (**B**) ADSCs CD34 Sca-1 positivity at 0, 48, and 72 hrs of culture. (**C**) MFI of CD24 in $CD34^{+}$ $Sca\text{-}1^{+}$ population at 0, 48, and 72 hrs.

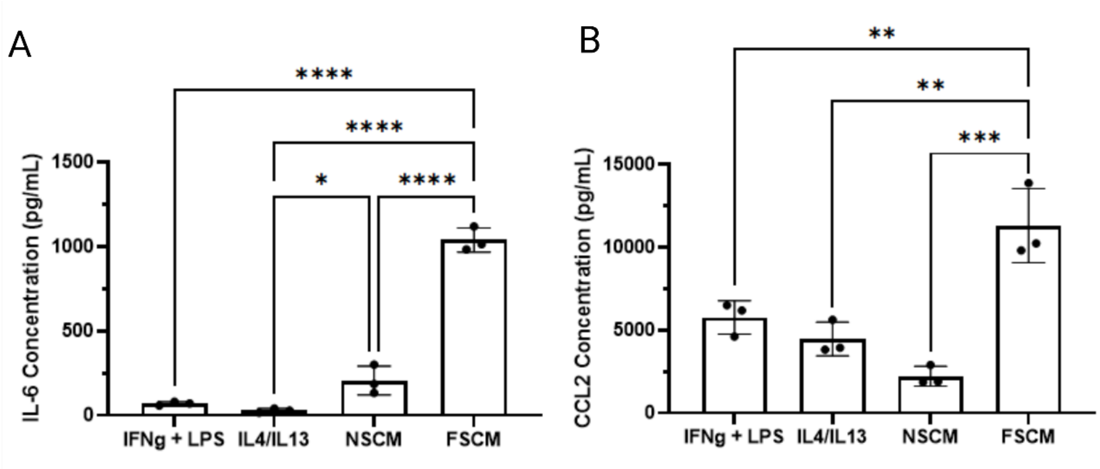


**Figure S3. CCL2 and IL-6 secretion after 48 hrs of co-culture.** BMDMs were activated with LPS/IFN-γ; IL-4/IL-13; normal skin conditioned media (NSCM) or fibrotic skin conditioned media (FSCM) and co-cultured with ADSCs for 48 hrs. Secreted levels of (**A**) IL-6 and (**B**) CCL2 in co-culture supernatants were measured by ELISA.

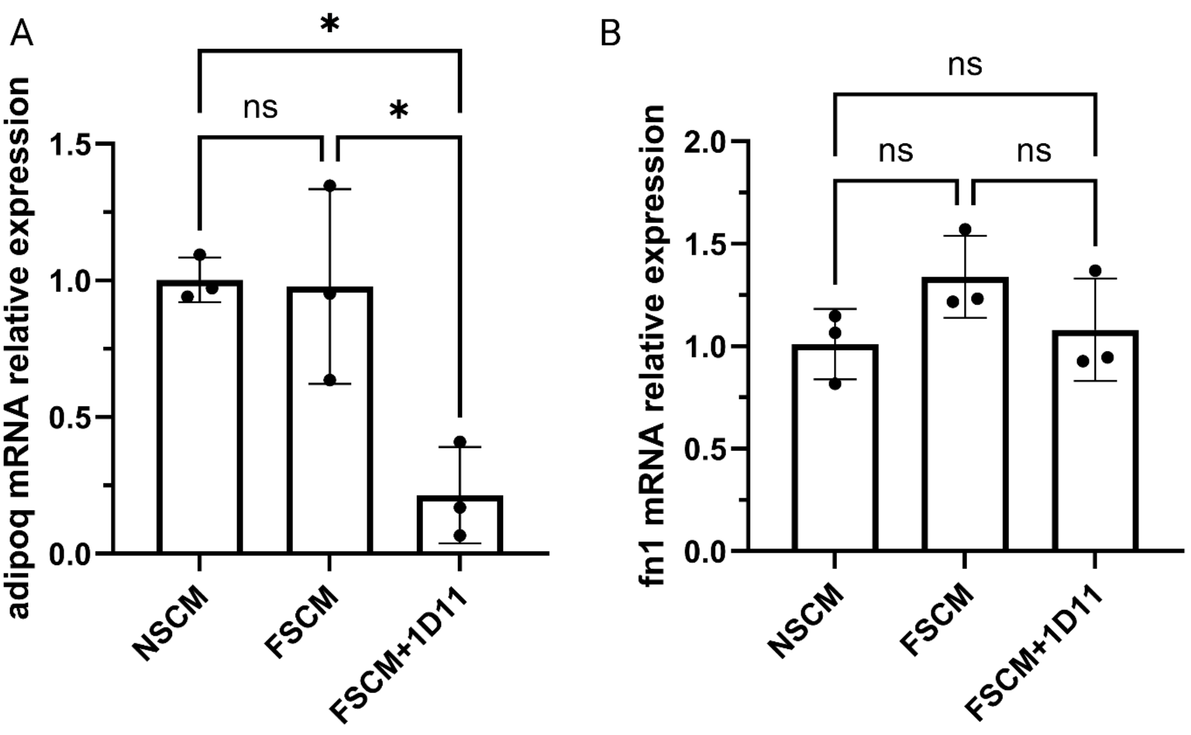


.

**Figure S4. qRT-PCR data from pan TGF-β blockade.** Relative mRNA levels of (**A**) ADIPOQ and (B) FN1 were assessed by qRT-PCR in differentiated ADSCs co-cultured with activated BMDMs for 96 hrs in the presence or absence of pan TGF-β blocking 1D11 antibody (2 μg/ml).

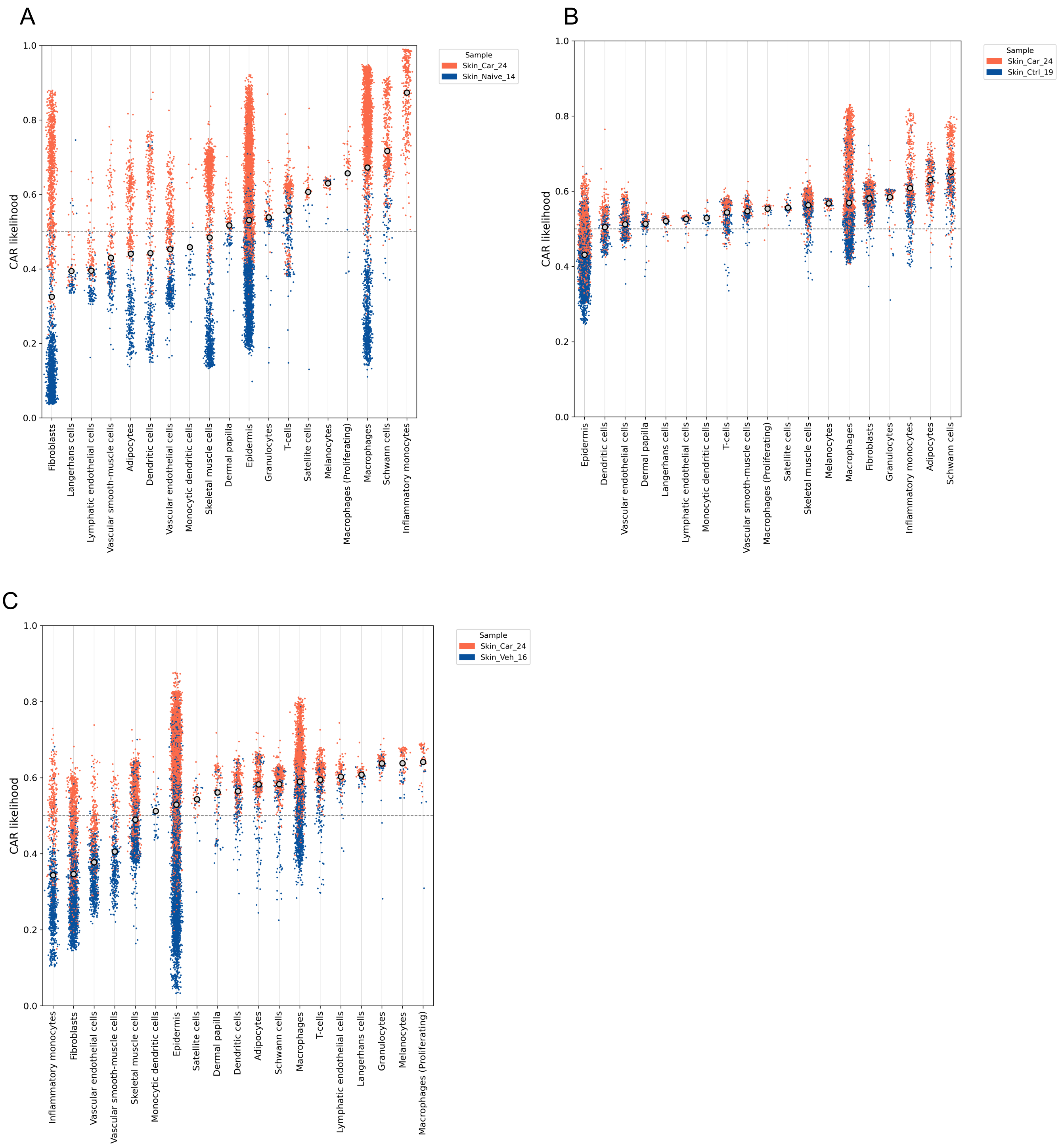


**Figure S5: Single-cell experimental perturbation analysis.** Anti-CD206 CAR T cell-associated relative likelihood values from experimental perturbation analysis of all identified cell types. Relative likelihoods quantify effect of treatment with anti-CD206 CAR T cells vs. reference groups (A: Naïve, B: BLM+HBSS, C: BLM+Control CAR T cells). Cells with likelihood values >0.5 are more likely to be found in the anti-

CD206 CAR T cell-treated sample, whereas cells with values <0.5 are more likely in the reference group. Color of points indicates sample of origin for each cell.
